# Kin-discriminating partner choice promotes the evolution of helping

**DOI:** 10.1101/2024.08.15.608103

**Authors:** Thomas W. Scott, Geoff Wild, Andy Gardner

## Abstract

Kin selection theory predicts that individuals should evolve to help relatives, either by helping indiscriminately in a population where they do not move very far from their relatives, or by discriminating kin and conditionally helping them. It has been argued that, because kin discrimination enables individuals to reduce how helpful they are with some social partners as well increase how helpful they are with others, this could lead to an increase or a decrease in the overall level of helping, depending on the curvature of the function relating the optimal level of help and genetic relatedness. However, this prediction was based on a model in which individuals were not able to choose their social partners but only adjust how helpful they should be towards those social partners they have been allocated. Here, we perform a mathematical analysis showing that if kin discriminators are allowed to choose whom they help, kin discrimination is more likely to increase the overall level of helping than previously anticipated. We obtained these results in two complementary theoretical settings: one more general and the other more specific and concrete.

## Introduction

Kin selection theory predicts that—at all levels of biology, from bacteria to humans— high genetic relatedness between social partners favours the evolution of helping (Hamilton, 1963, 1964; Maynard Smith, 1964; West *et al*., 2007). Genetic relatedness promotes helping because, by increasing a relative’s reproduction, a helpful individual is still passing its genes to the next generation, just indirectly. In viscous populations, where individuals do not disperse very far from their relatives, social partners tend to be closely related, such that individuals may evolve to help whomever they meet; this is called “indiscriminate helping” (Hamilton, 1964). Alternatively, in well-mixed populations, individuals may evolve to identify their relatives using either genetic or environmental cues, and then help them on a conditional basis; this is called “kin discrimination” (Green et al., 2024).

It is unclear whether overall levels of helping are expected to be higher or lower in indiscriminately helping versus kin-discriminating populations. Faria & Gardner (2020) pointed out that kin discriminators will evolve to help less when encountering nonrelatives as well as helping more when encountering close relatives, whereas indiscriminate helpers will exhibit an intermediate extent of helping with everyone. This difference between kin discriminators and non-discriminators implies there exists a geometric rule for determining which of these types displays the greater overall level of helping. If the optimal level of helping increases linearly with relatedness, indiscriminate helpers and kin discriminators are predicted to invest the same overall amount in helping. However, if the optimal level of helping is an accelerating (i.e. convex) function of relatedness, kin discriminators will disproportionately increase their help when encountering close relatives, resulting in more helping overall than compared with indiscriminate helpers. Conversely, if the optimal level of helping is a decelerating (i.e. concave) function of relatedness, kin discriminators will disproportionately decrease their help when encountering nonrelatives, resulting in less helping overall than compared with indiscriminate helpers.

Overall, Faria & Gardner (2020) argued that kin discrimination sometimes increases and sometimes decreases the overall level of helping. However, Faria and Gardner’s analysis did not account for an important characteristic of kin discrimination behaviour: while they assumed that kin discriminators can use relatedness to inform the degree to which they help a given partner, they neglected the possibility that discriminators might also use relatedness to inform partner choice (Scott et al., 2022, 2023; Scott, 2024). Partner choice may allow kin discriminators to reliably pick out close relatives, whom they will help intensively, increasing the overall level of helping more often, and to a greater degree, than anticipated by Faria and Gardner (2020).

We investigate this possible helpfulness-promoting effect of partner choice by analysing the evolution of cooperation to see when kin discrimination results in a higher overall level of helping than indiscriminate helping. First, we generalise Faria and Gardner’s (2020) analysis, allowing kin discriminators to engage in kin-discriminating partner choice as well as—or instead of—kin-discriminating helping. This reveals that kin-discriminating partner choice tends to promote helping, such that the overall level of help is higher under kin discrimination not only for linearity and convexity but also—in contrast to Faria & Gardner’s prediction—in some concavity scenarios. Second, we illustrate our general results by developing a concrete model of pairwise social interactions in a subdivided population, in which kin discriminators have multiple social encounters before deciding with whom to cooperate. We discuss the challenge that these results pose for the application of “veil of ignorance” thinking to social evolution.

### General analysis

We consider a population in which every individual enacts help towards one social partner. Each individual obtains its social partner in one of two ways. First, the individual samples *s* individuals from its neighbourhood. Then, in the “no kin-discriminating partner choice” scenario, a random individual from this sample is assigned as the individual’s social partner; we denote the individual’s relatedness to this social partner by *R*, and the expectation of this random variable by E(*R*). In the “kin-discriminating partner choice” scenario, the most highly related individual in the sample is assigned as the individual’s social partner; we denote the individual’s relatedness to this social partner by *R*_max_, and the expectation of this random variable by E(*R*_max_). On these assumptions, in the “no kin-discriminating partner choice” scenario, each individual is simply assigned a social partner. But in the “kin-discriminating partner choice” scenario, every individual has some choice as to who their social partner will be, with their choice being informed by their degree of relatedness to each potential social partner. In Appendix A we justify our assumption that individuals capable of kin-discriminating partner choice will choose the closest relative amongst the *s* possible social partners (rather than, for instance, the least related individual), by showing that this maximises their inclusive fitness. For greater *s*, individuals capable of kin-discriminating partner choice have a larger sample of potential social partners to choose from, meaning they are more likely to acquire a close relative, such that kin-discriminating partner choice is more effective.

How much help an individual enacts towards their social partner is also determined in one of two ways: in the “no kin-discriminating helping” scenario, individuals do not adjust their level of helping according to the relatedness of their social partner, and in the “kin-discriminating helping” scenario, individuals may adjust their level of helping according to the relatedness of their social partner. In the former scenario, individuals effectively commit to a level of helping before they have found a partner; in the latter scenario, the commitment comes after the partnership is set.

Accordingly, we consider four different scenarios by combining how partnerships and cooperation are determined in different ways: no kin-discriminating partner choice and no kin-discriminating helping (“N”); no kin-discriminating partner choice and kin-discriminating helping (“H”); kin-discriminating partner choice and no kin-discriminating helping (“PC”); and kin-discriminating partner choice and kin-discriminating helping (“PC+H”). The first scenario (N) corresponds to indiscriminate helping, and the latter three scenarios (H, PC and PC+H) correspond to different types of kin discrimination.

In the “no kin-discriminating helping” scenarios (N and PC), all individuals in the population are expected to exhibit the same level of helping; specifically, that which is optimised according to the expected relatedness of social partners. We define this as *z*_N_ = *Z*(E(*R*)) in the “no kin-discriminating partner choice” scenario (N) and *z*_PC_ = *Z*(E(*R*_max_)) in the “kin-discriminating partner choice” scenario (PC). In the “kin discriminating helping” scenarios (H and PC+H), individuals across the population are expected to vary in their levels of helping; specifically, they are expected to exhibit a level of helping which is optimised according to their specific relatedness to social partners. Accordingly, we define the population-average level of helping as *z*_H_ = E(*Z*(*R*)) in the “no kin-discriminating partner choice” scenario (H) and *z*_PC+H_ = E(*Z*(*R*_max_)) in the “kin-discriminating partner choice” scenario (PC+H). We assume that *Z* is an increasing function of its argument (relatedness). All details are summarised in Table 1. In Appendix B, we obtain the following results.

**Table 1.** Relatedness and overall level of helping in each of the four scenarios.

|  |  | Kin-discriminating partner choice? |  |  |  |
| --- | --- | --- | --- | --- | --- |
|  |  | No |  | Yes |  |
|  |  | Relatedness | Help | Relatedness | Help |
| Kin-discriminating helping? | No | $E(R)$<br>(partner not known when deciding level of help) | $z_N = Z(E(R))$ | $E(R_{\max})$<br>(partner not known when deciding level of help) | $z_{PC} = Z(E(R_{\max}))$ |
| | Yes | $R$<br>(partner known when deciding level of help) | $z_H = E(Z(R))$ | $R_{\max}$<br>(partner known when deciding level of help) | $z_{PC+H} = E(Z(R_{\max}))$ |

If the optimal level of helping (*Z*) is a linear function of its argument (relatedness), kin-discriminating helping neither increases nor decreases the overall level of helping (*z*_N_ = *z*_H_ and *z*_PC_ = *z*_PC+H_, owing to the linearity of *Z*), and kin-discriminating partner choice increases the overall level of helping (*z*_N_, *z*_H_ ≤ *z*_PC_, *z*_PC+H_, owing to *Z* being an increasing function of its argument; Figure 1b).

**Figure 1.**
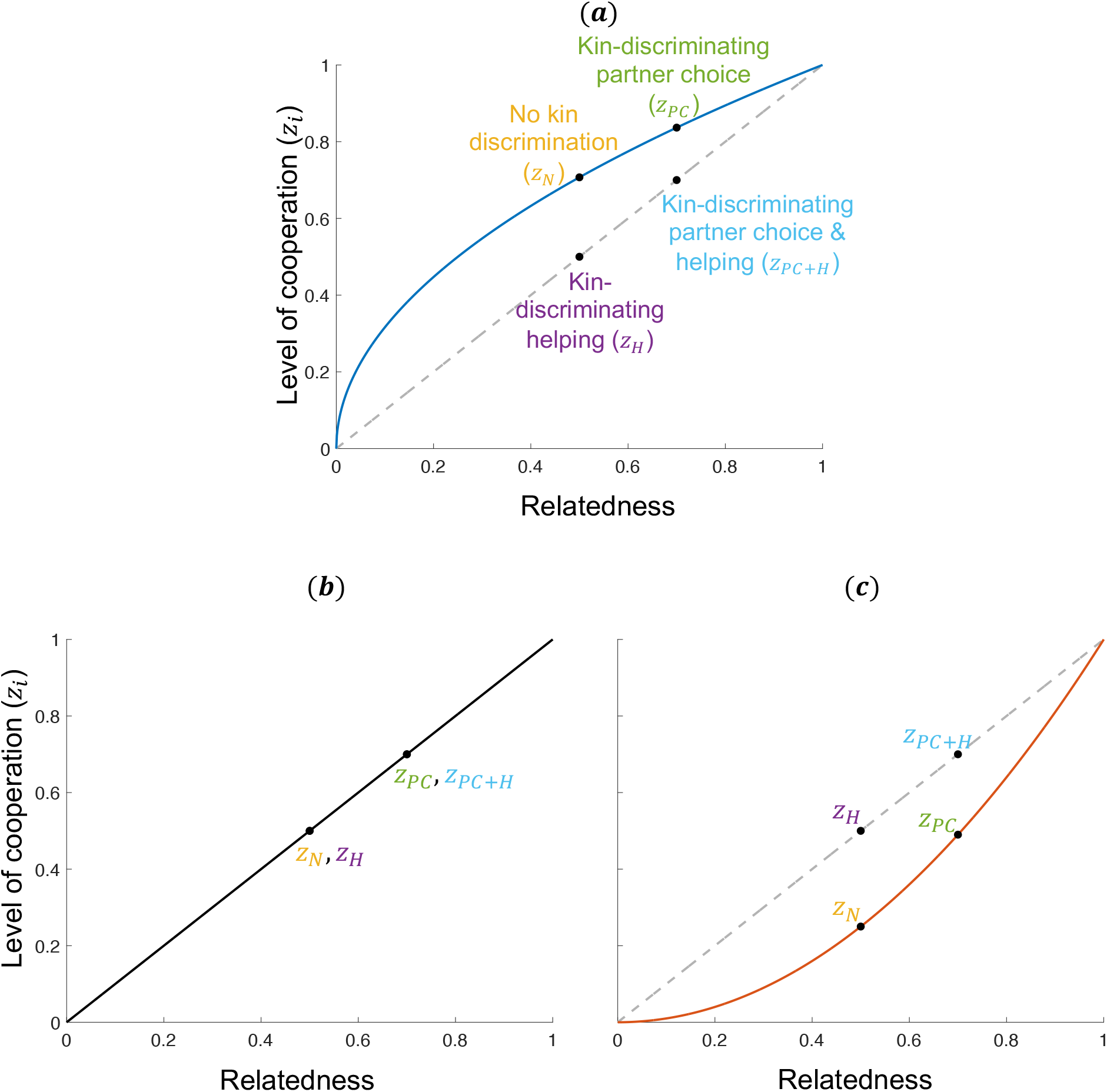
The effect of kin-discriminating helping and kin-discriminating partner choice on the overall level of helping. The black dots show the overall level of helping in four scenarios: no kin discrimination (*z*_N_); kin-discriminating helping (*z*_H_); kin-discriminating partner choice (*z*_PC_); kin-discriminating partner choice & helping (*z*_PC+H_). The solid lines in panels a–c show different relationships between the optimal level of helping and relatedness: concave (*a*); linear (*b*); convex (*c*). Kin-discriminating partner choice increases relatedness, shifting the dots rightwards along the solid lines, resulting in an increase in the overall level of helping (*z*_PC_ ≥ *z*_N_ & *z*_PC+H_ ≥ *z*_H_). Kin-discriminating helping pulls the overall level of helping away from the solid lines and onto the secants (grey dotted lines), which may increase (*c*; *z*_H_ ≥ *z*_N_ & *z*_PC+H_ ≥ *z*_PC_), decrease (*a*; *z*_H_≤ *z*_N_ & *z*_PC+H_ ≤ *z*_PC_) or leave unchanged (*b*; *z*_H_ = *z*_N_ & *z*_PC+H_ = *z*_PC_) the overall level of helping.

If the optimal level of helping (*Z*) is a concave (decelerating) function of its argument (relatedness), we find that kin-discriminating helping decreases the overall level of helping (*z*_N_ ≥ *z*_H_ and *z*_PC_ ≥ *z*_PC+H_, owing to the concavity of *Z*; Jensen, 1906; Faria & Gardner, 2020), and that kin-discriminating partner choice increases the overall level of helping (*z*_PC_ ≥ *z*_N_ and *z*_PC+H_ ≥ *z*_H_, owing to *Z* being an increasing function of its argument; Figure 1a). We also obtain the following subsidiary results. First, kin-discriminating partner choice without kin-discriminating helping increases the overall level of helping relative to kin-discriminating helping without kin-discriminating partner choice (*z*_PC_ ≥ *z*_H_, as implied by *z*_PC_ ≥ *z*_N_ and *z*_N_ ≥ *z*_H_). Second, kin-discriminating partner choice with kin-discriminating helping may or may not increase the overall level of helping relative to indiscriminate helping (*z*_N_ ≥ *z*_PC+H_ and *z*_N_ ≤ *z*_PC+H_ are both possible). That is, it could be that the helpfulness-boosting effect of kin-discriminating partner choice outweighs the helpfulness-inhibiting effect of kin-discriminating helping, or alternatively the reverse could be true.

If the optimal level of helping (*Z*) is a convex (accelerating) function of its argument (relatedness), we find that kin-discriminating helping increases the overall level of helping (*z*_N_ ≤ *z*_H_ and *z*_PC_ ≤ *z*_PC+H_, owing to the convexity of *Z*; Jensen, 1906; Faria & Gardner, 2020) and kin-discriminating partner choice also increases the overall level of helping (*z*_PC_ ≥ *z*_N_ and *z*_PC+H_ ≥ *z*_H_, owing to *Z* being an increasing function of its argument; Figure 1c). We also obtain the following subsidiary results. Firstly, kin-discriminating partner choice with kin-discriminating helping increases the overall level of helping relative to indiscriminate helping (*z*_PC+H_ ≥ *z*_N_, as implied by *z*_PC_ ≤ *z*_PC+H_ and *z*_PC_ ≥ *z*_N_). Secondly, kin-discriminating partner choice without kin-discriminating helping may or may not increase the overall level of helping relative to kin-discriminating helping without kin-discriminating partner choice (*z*_H_ ≥ *z*_PC_ and *z*_H_ ≤ *z*_PC_ are both possible). That is, it could be that the helpfulness-boosting effect of kin-discriminating partner choice exceeds the helpfulness-boosting effect of kin-discriminating helping, or alternatively the reverse could be true.

To sum up, although kin-discriminating helping *per se* might or might not boost overall levels of helping (with this depending on whether *Z* is convex or concave with respect to relatedness), kin-discriminating partner choice *per se* is expected to do so. If the two capacities always co-occur, then “kin discrimination” might or might not serve to boost overall levels of helping (*z*_PC+H_ ≥ *z*_N_ and *z*_PC+H_ ≤ *z*_N_ are both possible), but the dividing line does not coincide with the concavity / convexity distinction, *contra* Faria & Gardner (2020) (Figure 2). If *Z* is linear or convex (accelerating), then we expect “kin discrimination” to boost the overall level of helping (*z*_PC+H_ ≥ *z*_N_), but if *Z* is concave (decelerating) then “kin discrimination” has the potential to increase or decrease the overall level of helping (*z*_PC+H_ ≥ *z*_N_ and *z*_PC+H_ ≤ *z*_N_ are both possible). We provide a graphical interpretation of these results in Figures 1 & 2.

**Figure 2.**
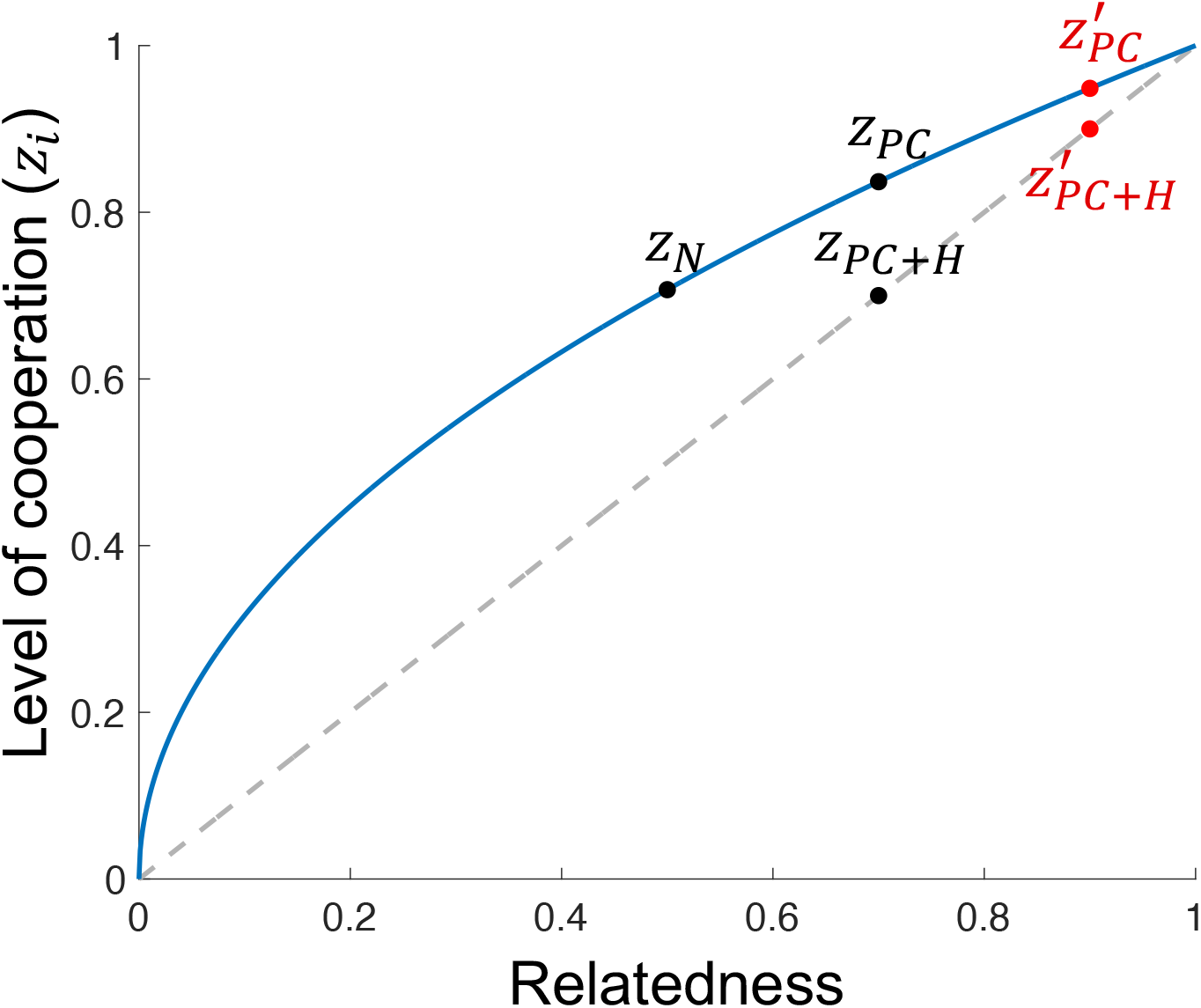
Kin discrimination increases the overall level of helping with sufficient partner choice, but need not in general. The black dots show the overall level of helping when there is: no kin discrimination (*z*_N_); kin-discriminating partner choice (*z*_PC_); kin-discriminating partner choice & helping (*z*_PC+H_). The red dots show the impact of further increasing partner choice, s 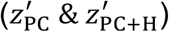. The solid line represents the relationship between the optimal level of helping and relatedness, which we assume here is concave. Kin-discriminating partner choice increases relatedness, which shifts the dots rightwards along the solid lines, increasing the overall level of helping; kin-discriminating helping pulls the overall level of helping away from the solid line and onto the secant, decreasing the overall level of helping. There are two key results: (1) if the capacities for kin-discriminating helping and partner choice coincide, this may decrease the overall level of helping (*z*_PC+H_ < *z*_N_) or increase it 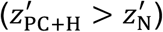, relative to indiscriminate helping; (2) for increased kin-discriminating partner choice, s, the influence of kin-discriminating helping on the overall level of helping, mediated by the curvature of the solid line, is reduced, as is evidenced by 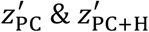 being closer to each other than *z*_PC_& *z*_PC+H_.

The reason why kin-discriminating partner choice *per se* consistently increases the overall level of helping is that it allows individuals to pick out closer relatives, who elicit a higher level of helping. The reason why kin-discriminating helping *per se* does not necessarily increase the overall level of helping is that it simply allows individuals to fine-tune their level of helping to the relatedness of each social partner, but there is no guarantee that this fine tuning will result in greater helping overall (Jensen, 1906; Faria & Gardner, 2020).

In Appendix C we obtain a further result, which is that, as partner choice becomes more effective (i.e., as *s* is increased), kin-discriminating helping has less effect on the overall level of helping, with no effect in the limit of perfect partner choice (Figure 2). This result arises because, with more effective partner choice, there is less relatedness variation amongst social partners, meaning there is no phenotypic consequence of being able to adjust one’s level of help towards a given social partner. The implication of this result is that, if the capacities of kin-discriminating helping and kin-discriminating partner choice coincide, kin discrimination will increase the overall level of helping if there is sufficient partner choice (i.e., if *s* is sufficiently high).

Note that if we relax the assumption that *Z* is increasing and, indeed, say that it is constant with respect to its argument (relatedness), we find that there is no impact of kin-discriminating helping or kin-discriminating partner choice on the overall level of helping (*z*_N_ = *z*_H_ = *z*_PC_ = *z*_PC+H_). In this case, we would still expect individuals to choose high-relatedness social partners, as it benefits one’s inclusive fitness to provide a standard unit of help to a closer relative than to a more distant relative (Appendix A). This would apply to empirical scenarios where individuals distribute resources of fixed size to other individuals. These resources could be nests, territories, or even the actor’s own body as a food offering as in instances of matriphagy. This contrasts with the more general scenario, in which individuals can decide how much help to give, as well as to whom it is given.

### Illustration

We illustrate the results of our analysis in more concrete terms, by developing a model of pairwise social interactions in a subdivided population, where kin discriminators are allowed to have multiple social encounters before deciding who to help. We assume a patch-structured population with *n* ≥ 2 hermaphrodite diploid breeders in each patch. Each breeder produces the same large number of offspring before dying, and these offspring interact socially within their patch, in a way that affects their survival to adulthood. All surviving individuals disperse away to seek reproductive opportunities elsewhere, mating monogamously with unrelated mating partners *en route* and arriving at a new patch. Following dispersal, *n* individuals are chosen at random on each patch to be breeders and all other individuals perish, returning the population to the beginning of the lifecycle.

These assumptions mean that each juvenile individual is related to a proportion 1/*n* of her patch-mates by one-half (siblings) and a proportion (*n*-1)/*n* by zero (nonrelatives) (Michod & Hamilton, 1980). We assume that, in the lifecycle stage where social interactions occur, each individual obtains a ‘recipient’ by choosing the most-related individual from a sample of *s* of her patch mates, chosen randomly with replacement (Scott et al., 2022, 2023; Scott, 2024). Consequently, each individual has a probability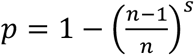 of obtaining a sibling rather than a nonrelative recipient. This yields *p* = 1/*n* for *s* = 1 (no kin-discriminating partner choice) and *p* → 1 as *s* → ∞ (perfect kin-discriminating partner choice).

Individuals then have the opportunity to help their recipients. A focal individual invests *z* into helping, at a personal survival cost *C*(*z*), to give a survival benefit *B*(*z*) to her recipient. We assume the following functional forms: 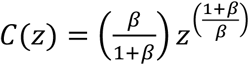 and *B*(*z*) = *z*, where 0 < β < ∞, such that the focal individual’s inclusive fitness - *C*(*z*) + *B*(*z*) *r* is maximized at

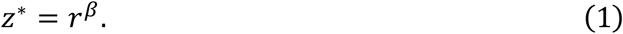

Inspecting Equation (1) reveals that the *β* parameter determines the curvature of the function *Z*(*r*) = *r*^β^ relating the optimal level of helping and relatedness, with 0 < β < 1 yielding a concave relationship, *β* = 1 yielding a linear relationship and *β* > 1 yielding a convex relationship.

If there is neither kin-discriminating partner choice nor kin-discriminating helping (N) then a focal individual’s relatedness valuation of her recipient is *r* = E(*R*) = 1/(2*n*), and making this substitution into Equation (1) yields a level of helping

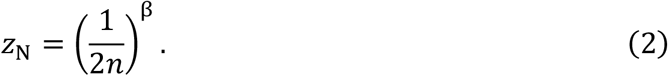

If there is no kin-discriminating partner choice but there is kin-discriminating helping (H) then a focal individual’s relatedness valuation of her recipient is *r* = *R*, i.e. with probability *p* = 1/*n* her recipient is a sibling and hence related by *R* = ½, such that from Equation (1) she helps by an amount *z*\* = (1/2) ^β^, and with probability 1-*p* = (*n*-1)/*n* her recipient is a non-relative and hence related by *R* = 0, such that from Equation (1) she helps by an amount *z*\* = 0. Taking an average over all actors, this yields an overall level of helping

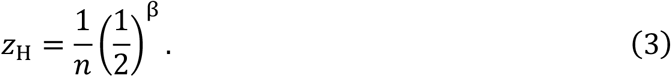

If there is kin-discriminating partner choice but no kin-discriminating helping (PC) then a focal individual’s relatedness valuation of her recipient is *r* = E(*R*_max_), i.e. with probability 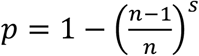 her recipient is a sibling and hence related by *R*_max_ = ½ and with probability 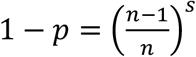 her recipient is a non-relative and hence related by *R*_max_ = 0, which yields an expected relatedness of 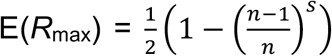 Making this substitution into Equation (1) yields an overall level of helping

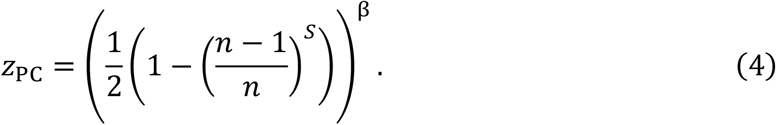

Finally, if there is both kin-discriminating partner choice and kin-discriminating helping (PC+H) then a focal individual’s relatedness valuation of her recipient is *r* = *R*_max_, i.e. with probability 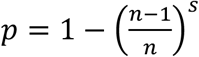 her recipient is a sibling and hence related by *R*_max_ = ½, which from Equation (1) elicits a level of helping *z*\* = (1/2)^β^, and with probability 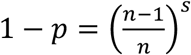 her recipient is a non-relative, which from Equation (1) elicits a level of helping *z*\* = 0. Taking an average over all actors, this yields an overall level of helping

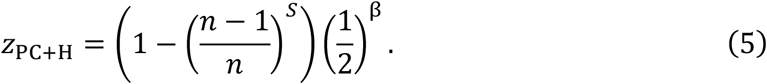

Comparing our *z* values reveals that: if 0 < β < 1 (i.e. concavity) then we have *z*_N_ ≥ *z*_H_, *z*_PC_ ≥ *z*_PC+H_, *z*_PC_ ≥ *z*_N_ and *z*_PC+H_ ≥ *z*_H_; if *β* = 1, (i.e. linearity) then we have *z*_N_ = *z*_H_ ≤ *Z*_pc_ = *z*_PC+H_; and if *β* > 1 (i.e. convexity) then we have *z*_N_ ≤ *z*_H_, *z*_PC_ ≤ *z*_PC+H_, *z*_PC_ ≥ *z*_N_, *z*_PC+H_ ≥ *z*_H_. We also find that *z*_PC+H →_ *z*_PC_ as *s* → ∞. That is, this illustrative treatment exactly recovers the results of our general analysis. In Appendix D we undertake a more formal “neighbour-modulated fitness” analysis of the illustrative model and obtain identical results to the more intuitive inclusive fitness analysis presented here (Taylor et al., 2007; Scott & Wild, 2023).

## Discussion

Kin discrimination can involve one or both of the following: (1) kin-discriminating partner choice, in which individuals choose who to help; (2) kin-discriminating helping, in which individuals choose how much to help a given partner. Previous theory developed by Faria & Gardner (2020) on the consequences of kin discrimination for the overall level of helping has considered (2) but not (1). We found that kin-discriminating partner choice allows individuals to pick out closer relatives, who elicit a higher level of helping, increasing the overall level of helping relative to when individuals help indiscriminately (Figure 1). In contrast, kin-discriminating helping allows individuals to fine-tune their level of helping to the relatedness of each social partner, and this fine tuning may increase (Figure 1c), decrease (Figure 1a) or leave unchanged (Figure 1b) the overall level of helping relative to when individuals help indiscriminately (Faria & Gardner, 2020). Therefore, if the capacities for kin-discriminating partner choice and helping coincide, then kin discrimination might or might not serve to boost the overall level of helping, but will do so if there is sufficient kin-discriminating partner choice (Figure 2).

It has been argued that a “veil of ignorance” tends to have a promoting effect on the evolution of helping. In the present context the idea—which has been imported to evolutionary biology from moral philosophy (Harsanyi, 1955; Rawls J, 1971; Frank, 2003; Okasha, 2012)—is that if individuals know how related they are to their social partners they have the opportunity to restrict help to nonrelatives, but if individuals are ignorant with respect to the relatedness of their social partners they cannot identify their nonrelatives and so do not have the opportunity to restrict help. Consequently, ignorant individuals (indiscriminate helpers) should help more than knowledgeable individuals (kin discriminators) (Queller & Strassmann, 2013). For example, in social insects where the queen mates multiple times, colonies comprise different classes of relative (patrilines) and individuals usually can’t tell which nest-mates are more closely related (Keller, 1997; Queller & Strassmann, 2002). It has been assumed that this ignorance promotes cooperation within the colony (Queller & Strassmann, 2013).

This veil-of-ignorance thinking in evolutionary biology has recently been challenged by Faria & Gardner (2020). They pointed out that, although ignorant individuals do not have the opportunity afforded to informed individuals of identifying and restricting help towards nonrelatives, they also lack the ability to identify and upregulate help towards relatives. Consequently, a veil of ignorance will only increase the overall level of helping if the informed downregulation of help towards nonrelatives is greater than the informed upregulation of help towards relatives (Faria & Gardner, 2020). Faria & Gardner (2020) showed that this will only be the case if the optimal level of helping is a concave (*i*.*e*. decelerating) function of relatedness. Consequently, for the veil-of-ignorance argument to be convincing, we would also need an account of why helping should increase concavely with relatedness; no such account has yet been given.

Our work provides a further challenge to veil-of-ignorance thinking in evolutionary biology. In nature, individuals who are informed as to their relatedness to social partners (kin discriminators) are often able to choose their social partners, unlike ignorant individuals (indiscriminate helpers). Consequently, informed individuals choose close relatives, with whom they help intensively, tending to increase the overall level of helping relative to ignorant individuals, in opposition to veil-of-ignorance thinking. The reason for this discrepancy is that, in nature but not moral philosophy, lifting the veil of ignorance does not just reveal to individuals their social partners; it also reveals the identities of other potential social partners, whom the individuals may choose to help instead. In other words, in nature, kin discriminators and indiscriminate helpers differ from each other not only with regard to their ignorance over relatedness, but also with regard to whether they can choose their social partners. This extra biological detail (partner choice) means that veil-of-ignorance thinking in moral philosophy is not always directly analogous to veil-of-ignorance thinking in evolutionary biology, so may not tend to work in the same way. A broader point here is that care should be taken when applying theories developed in the social sciences or philosophy to evolutionary biology, as biological complexities may weaken the analogy and lead to opposite predictions; formal modelling can be useful here (Scott & West, 2019). We note that veil-of-ignorance thinking may still be important in biology, for instance as a means of reducing within-group conflict, even if the overall helping investment is not typically increased (Queller & Strassmann, 2013; Rautiala & Gardner, 2023).

Our work emphasises the importance of kin-discriminating partner choice in promoting helpfulness. This relates to recent theoretical work showing that partner choice is often required for kin discrimination to evolve in the first place (Scott et al., 2022, 2023; Scott, 2024). Putting these two strands of research together, we might therefore expect that, when we do observe kin discrimination in nature, partner choice is likely to also be present (as a pre-requisite), and therefore kin discrimination is likely to be promoting the level of helpfulness. It would be very useful to test these theoretical predictions empirically. Partner choice appears to be very common in kin discriminating animals, such as cooperatively breeding birds and mammals, in which individuals can move around and encounter multiple potential social partners within their group or larger social network before deciding who to help (Scott et al., 2022; Green et al., 2024).

## Acknowledgements

For funding, we thank the ERC (834164 & 771387), NERC (NE/V011537/1) and the Natural Sciences and Engineering Research Council of Canada (RGPIN-2019-06626).

## Appendix A

We show in this appendix that kin-discriminating partner choice causes individuals to choose partners with higher (rather than lower) relatedness. We assume that an individual has a choice of social partners with varying relatedness (*r*), and then enacts an amount of helping that is optimal according to the relatedness of the partner they have chosen. The inclusive fitness of the individual (actor) can be written as

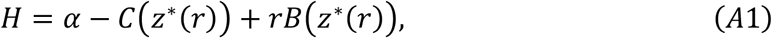

where: *α* is baseline fitness; *C*(*z*) is the cost of helping by an amount *z*; *B*(*z*) is the benefit of receiving an amount of help *z*; *r* is the relatedness of actor and recipient; *z*\*(*r*) is the optimal level of help as a function of relatedness.

Then, if the individual can choose to effectively increase the relatedness of the social partner to whom she directs her help, she would experience a marginal change in inclusive fitness given by:

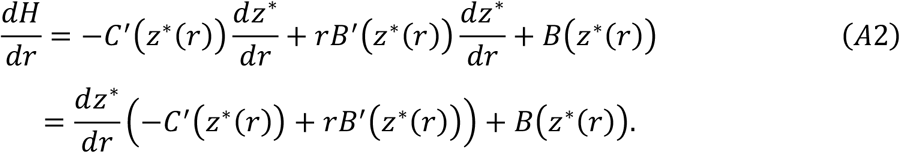

By virtue of *z*^∗^(*r*) being the optimal level of help, it satisfies the condition −*C*′(*z*^∗^(*r*)) + *rB*′(*z*^∗^(*r*)) = 0. This means that we can simplify Equation A2 to obtain

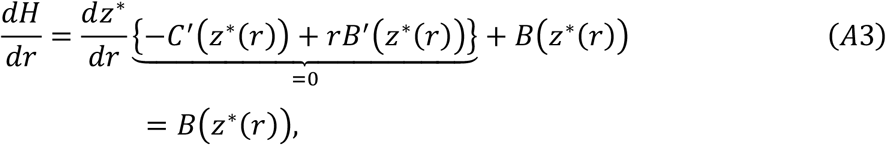

which is positive, so the individual’s inclusive fitness is always increased by choosing a social partner of higher relatedness. This means that, if an individual is allowed to encounter more than one individual before choosing her social partner (kin-discriminating partner choice), she will choose the closest relative.

## Appendix B

In this appendix, we obtain the results presented in the main text general analysis. An actor samples *s* individuals from her neighbourhood. Let *R*_i_ denote the relatedness between the actor and the *i*th individual sampled, let 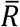 denote the mean of the sample, and let *R*_max_ be the largest value sampled. Obviously, 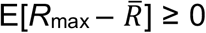, where E is an expectation taken with respect to the joint distribution of {*R*_1_, …, *R*_*s*_}. It follows that the expected value of *R*_max_, E[*R*_max_], is never less than the expected value of the sample mean, 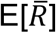. If {*R*_1_, … *R*_*s*_} is an iid sample from a distribution with expectation *μ*_*r*_, then we can go further and say that E[R_max_] is never less than *μ*_*r*_, because 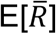 is an unbiased estimator of *μ*_*r*_ in this case.

Let *Z* be an increasing function of a single variable. Then we have 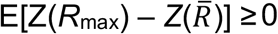, which we can re-write as z_PC+H_ ≥ z_H_ (in keeping with the main text). We also have 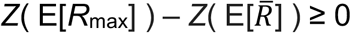,which we can re-write as z_PC_ ≥ z_N_ (in keeping with the main text). If *Z* is convex, then Jensen’s (1906) inequality says *Z*(E[*R*_max_]) ≤ E[*Z*(*R*_max_)] and 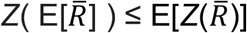. It follows that, for increasing and convex *Z*, we have

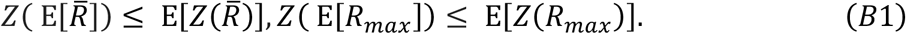

Thinking of *Z* as the optimal level of helping, we re-write the previous line as, *z*_N_ ≤ *z*_H_, *z*_PC_ ≤ *z*_PC+H_ (in keeping with the main text). When kin discrimination occurs, we evaluate *Z* before the expectation (the actor decides on a level of help after sampling); when it does not occur, we evaluate *Z* after the expectation (the actor decides on a level of help before sampling). When kin-discriminating partner choice occurs, we evaluate at *R*_max_ and when it does not occur we evaluate at 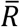.

If *Z* is concave then *Z*(E[*R*_max_]) ≥ E[*Z*(*R*_max_)] and 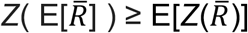, and we have

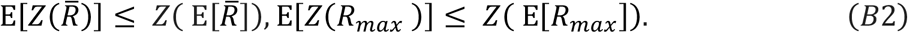

This can be re-written as *z*_H_ ≤ *z*_N_, *z*_PC+H_ ≤ *z*_PC_ (in keeping with the main text).

If *Z* is linear, then *Z*(E[*R*_max_]) = E[*Z*(*R*_max_)] and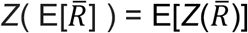, and we have

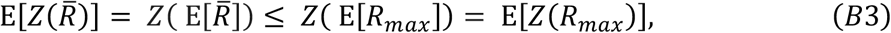

or simply, *z*_N_ = *z*_H_ ≤ *z*_PC_ = *z*_PC+H_ as in the main text.

## Appendix C

In this appendix we show that, as partner choice becomes more perfect, kin-discriminating helping has less effect on the overall level of helping, with no effect in the limit of perfect partner choice.

We collect a sample of *s* independent and identically distributed observations. Let *F*(*x*) be the cumulative distribution function associated with the probability distribution from which the observations have been taken. We will assume that there is a number *r*_1_ such that *F*(*r*) = 1 and *F*(*r*) < 1 for any *r* < *r*_1_. In other words, we will assume that there is a maximum attainable value, *r*_1_, that any given observation may achieve.

### Proposition 1.

*Let R*_*max*_ *be the largest observation in the sample. The expectation of R*_*max*_ *can be made arbitrarily close to r*_1_ *by choosing a sufficiently large sample size, s*.

*Proof*. The proof follows from combining two separate observations. We first observe that given any *ε* such that 0 < *ε* < *r*_1_, the quantity

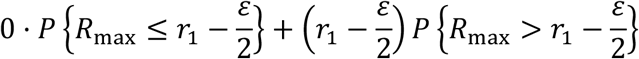

cannot overestimate the expectation of *R*_max_. In other words,

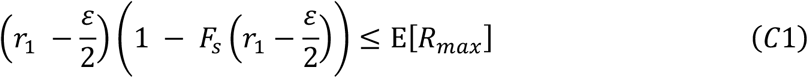

where *F*_*s*_(*r*_1_) is the cdf associated with *R*_max_. Second, we observe that the quantity 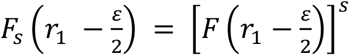 tends to zero – in a pointwise fashion – in the limit of large *s*. Thus, there exists an integer *s*_1_ such that

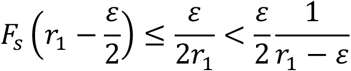

for all *s* > *s*_1_. It follows that

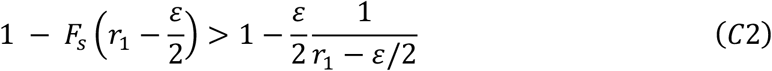

whenever *s* > *s*_1_. Combining inequalities (C1) and (C2), we conclude that for sufficiently large *s*,

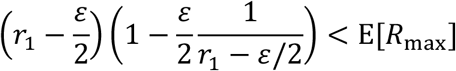

which simplifies to 0 < *r*_1_ − E[*R*_*max*_] < *ε*, as required. □

### Corollary to Proposition 1.

*For continuous* Z, *the quantity Z*(E[*R*_max_]) *can be made arbitrarily close to Z*(*r*_1_) *by choosing a sufficiently large sample size, s*.

We suppose that *Z*(*r*) is a strictly increasing function of *r*, with *Z*(0) = 0. We suppose further that *Z* is differentiable on (0, *r*_1_] with |*Z*′(*r*)| ≤ *M*, where it is understood that *Z*′(*r*_1_) is calculated using a limit from below. [Alternatively, we could consider a smooth extension of *Z* that expands the domain to include some *r*_2_ > *r*_1_. Then we would assume that, for the extended function, |*Z*(*r*)| ≤ *M* for all *r* in an open interval containing *r*_1_.] For later use, we note that for a sufficiently small *ε* > 0 we also have 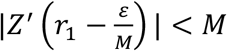. Moreover, from Taylor’s Remainder Theorem we have

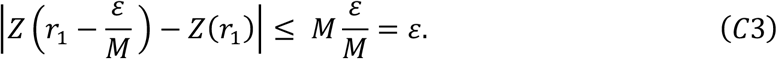

Given that *Z* is increasing the previous line simplifies to 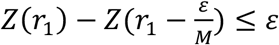 or

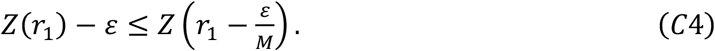

### Proposition 2.

*If Z has the properties described above, then the expectation of Z*(*R*_max_) *can be made arbitrarily close to Z*(*r*_1_) *by choosing a sufficiently large sample size, s*.

*Proof*. We treat (small) *ε* > 0 as given and observe that, because *Z* is a strictly increasing function, the quantity

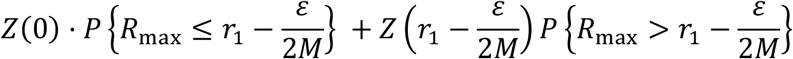

cannot overestimate the expectation of *Z*(*R*_max_). Using the fact that *Z*(0) = 0, this observation becomes

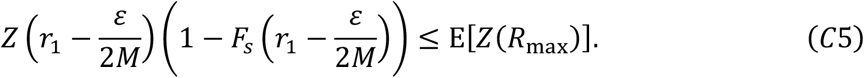

If we incorporate the information given in (C4) into (C5), we obtain

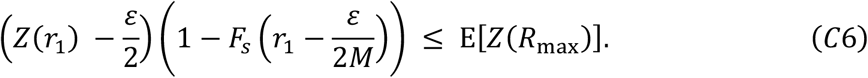

As we observed in the proof to Proposition 1, 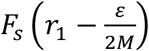 converges pointwise to 0 in the limit of large sample size *s*. Therefore we can find a sample size s that is large enough to guarantee

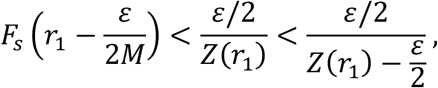

or equivalently

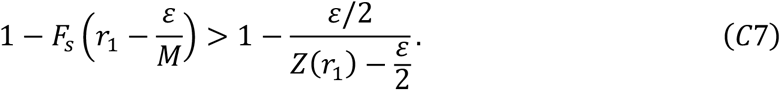

Combining inequalities (C6) and (C7), we obtain

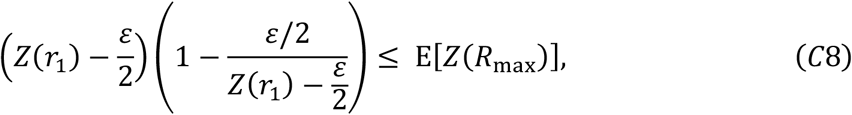

which simplifies to *Z*(*r*_1_) − *ε* ≤ E[*Z*(*R*_max_)], as required. □

## Appendix D

In this appendix, we analyse the illustrative model presented in the main text using a “neighbour-modulated fitness” rather than an “inclusive fitness” approach (Taylor et al., 2007; Scott & Wild, 2023). Our lifecycle assumptions mean that each individual is chosen to be someone’s recipient, in expectation, one time per generation. A focal individual therefore receives an expected overall helping investment of *y*_*ij*_, where *y*_*ij*_ is the average helping investment amongst individuals who choose the focal individual as their recipient. We denote the population average helping investment by *z*_*ij*_.

Consequently, the focal individual incurs a cost of helping *C*(*x*_*ij*_) and receives a benefit of helping *B*(*y*_*ij*_). We assume the following functional forms: 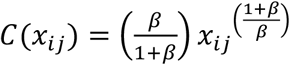 & *B*(*y*_*ij*_) = *y*_*ij*_.

The focal individual’s survival probability will be proportional to the following (recall that 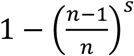 gives the probability of helping, or equivalently being helped by, a sibling, and 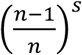 gives this probability for a non-sibling):

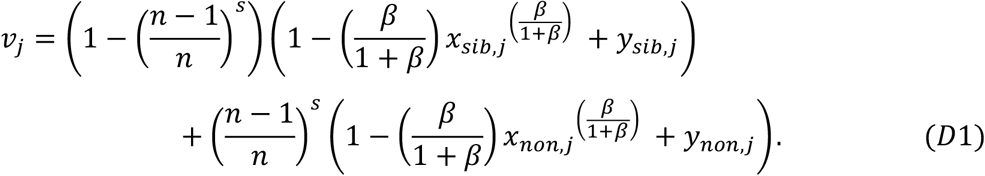

Substituting *z*_*ij*_ in for *x*_*ij*_ and *y*_*ij*_ in Equation D1, we can obtain the survival probability for a random individual drawn from the population. It is proportional to

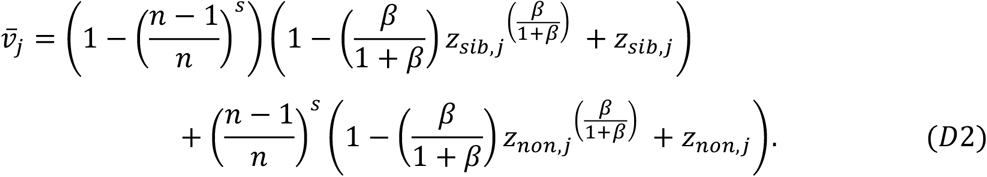

The focal individual’s neighbour-modulated fitness, *i*.*e*., her number of offspring after one full iteration of the lifecycle, is then obtained as

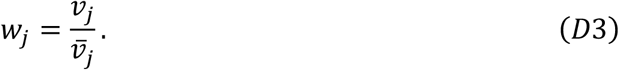

The equilibrium levels of helping enacted towards siblings 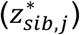 and nonrelatives 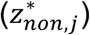 will satisfy the following conditions:

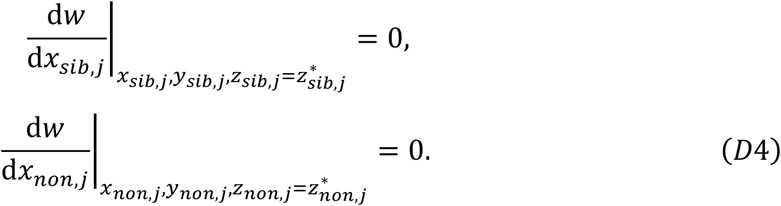

These conditions can be written equivalently as follows using the chain rule, where each derivative and partial derivative below is evaluated at the equilibrium 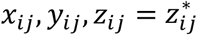:

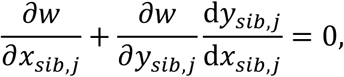

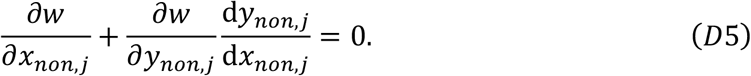

(Taylor & Frank, 1996; Frank, 1998; Rousset, 2004; Brown & Taylor, 2010; Lehmann & Rousset, 2010). To evaluate these conditions, we first note that 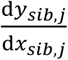 and 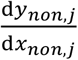 give the marginal change in partner trait value with a change in actor trait value, equivalent to the coefficient of relatedness between the actor and its social partner (*r*_*ij*_) (Taylor & Frank, 1996; Frank, 1998; Rousset, 2004). We can therefore evaluate 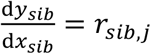 and 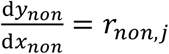. We obtain

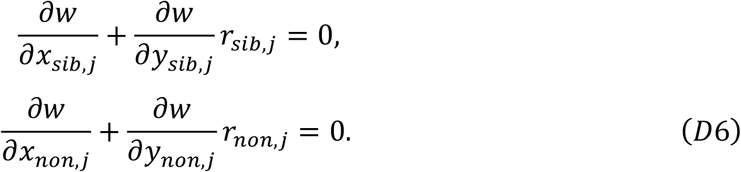

Next, we can obtain the partial derivatives 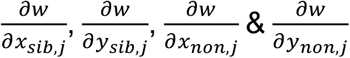 by differentiating Equation D3 with respect to *x*_*sib,j*_, *y*_*sib,j*_, *x*_*non,j*_ & *y*_*non,j*_ respectively. Evaluating everything at the equilibrium 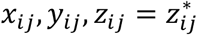, we obtain the equilibrium levels of helping towards siblings and nonrelatives, respectively, as

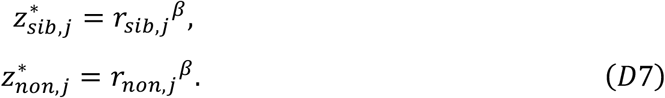

Therefore, the *β* parameter solely determines whether the optimal level of helping increases linearly (*β* = 1), convexly (*β* > 1) or concavely (*β* < 1) with relatedness.

In the “kin-discriminating helping” scenario (*j* = *disc*), individuals know how related they are to each of their social partners, and can adjust their level of help accordingly, such that 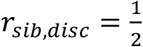 and *r*_*non,disc*_ = 0. Substituting these into Equation D7, we obtain the equilibrium levels of helping towards siblings and nonrelatives in the “kin-discriminating helping” scenario as

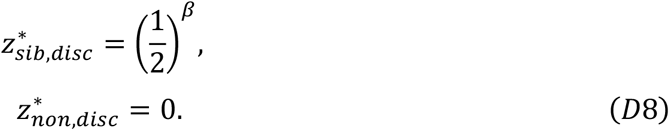

Helping nonrelatives is disfavoured 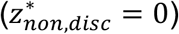 because it provides no indirect or direct fitness benefits to the actor.

The average level of helping in the “kin-discriminating helping” scenario, 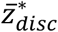, is calculated by summing the weighted levels of helping towards siblings and nonrelatives, where the weights are probabilities of interaction (*i*.*e*., by taking 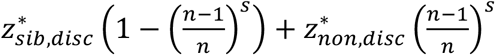. We obtain

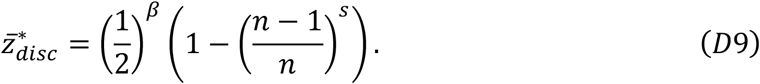

In the “no kin-discriminating helping” scenario (*j* = *indisc*), in which individuals help indiscriminately, individuals must commit to a given level of help before they have observed whether their social partner is a sibling or nonrelative. This level of help will be optimised with respect to the average relatedness between social partners, given by 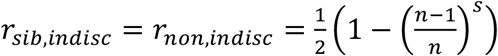. Substituting these into Equation D7, we obtain the equilibrium level of helping (towards siblings, nonrelatives, and on average) in the “no kin-discriminating helping” scenario as

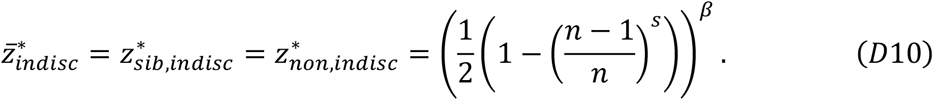

Note that, as *s* tends to infinity, 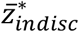 (Equation D10) and 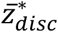 (Equation D9) converge on 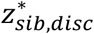 (Equation D8). That is, with perfect partner choice, kin discriminators pick out relatives with certainty, meaning the equilibrium level of helping becomes the optimal level of helping towards relatives, and this is true irrespective of whether individuals are capable of kin-discriminating helping.

First, we ask when kin-discriminating partner choice *per se* increases the overall level of helping (*i*.*e*., holding fixed the level of kin-discriminating helping). Kin-discriminating partner choice is encapsulated by the *s* parameter, with *s* =1 implying no kin-discriminating partner choice, and greater *s* implying greater kin-discriminating partner choice. Using Equations D9 & D10, we can observe that 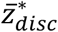and 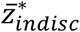are increasing functions of *s*. Therefore, kin-discriminating partner choice *per se* increases the overall level of helping in the population, and this is true irrespective of whether the population exhibits kin-discriminating helping.

Second, we ask when kin-discriminating helping *per se* increases the overall level of helping (*i*.*e*., holding fixed the level of kin-discriminating partner choice). We examine this by comparing the values of 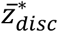 (Equation D9) and 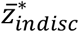 (Equation D10). We recover Faria & Gardner’s result, finding that: 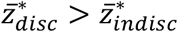 whenever 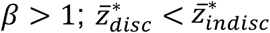 whenever 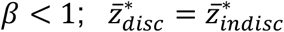 whenever *β* = 1. This implies that kin-discriminating helping: increases the overall level of helping when the optimal level of help increases convexly with relatedness (*β* > 1); decreases the overall level of helping when the optimal level of help increases concavely with relatedness (*β* < 1); neither increases nor decreases the overall level of helping when the optimal level of help increases linearly with relatedness (*β* = 1).

Finally, we ask, when the capacities for kin-discriminating helping and kin-discriminating partner choice coincide, when does this increase the overall level of helping? In other words, when is a population exhibiting both kin-discriminating helping and kin-discriminating partner choice more helpful than a population exhibiting neither? We examine this by evaluating 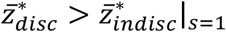, using Equations D9 and D10. We obtain the following condition for when kin discrimination increases the overall level of helping:

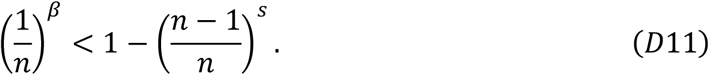

We can see that this condition is more likely to be satisfied for higher *β* and *s*. That is, kin discrimination is more likely to increase the overall level of helping when there is more partner choice and when the optimal level of help increases more convexly with relatedness. We can also see that this condition is not always satisfied. Specifically, kin-discriminating partner choice has a helpfulness-boosting effect, which increases with *s*. Kin-discriminating helping may have a helpfulness-boosting effect, which increases as *β* increases above 1, or a helpfulness-inhibiting effect, which increases as *β* decreases below 1. Kin discrimination only increases the overall level of helping if there is a net helpfulness-boosting effect.

We also highlight that, although kin-discriminating helping and kin-discriminating partner choice may both affect whether the overall level of helping is increased or decreased by kin discrimination, when these capacities coincide, the effect of kin-discriminating partner choice dominates. To see this, note that, as kin-discriminating partner choice (*s*) increases from 1 to infinity, the curvature of returns from helping (*β*) goes from fully determining whether Equation D11 is satisfied (*β* > 1) or not (*β* ≤ 1), to having no influence over it at all. This result makes sense because, for increased kin-discriminating partner choice, the relatedness between kin discriminators and their social partners becomes less variable (*i*.*e*., closer relatives are picked out more reliably). Consequently, for increased kin-discriminating partner choice, kin discriminators invest less variably in helping across social interactions (*i*.*e*., investment is high more reliably). Therefore, for increased kin-discriminating partner choice, the kin-discriminating helping component of kin discrimination becomes less phenotypically visible, minimising the effect of kin-discriminating helping on the overall level of helping.

